# Molecular Structure Study on the Polyelectrolyte Properties of Actin Filaments

**DOI:** 10.1101/2022.01.07.475417

**Authors:** Santiago Manrique-Bedoya, Marcelo Marucho

**Author notes:** Email address (Marcelo Marucho).

## Abstract

An accurate characterization of the polyelectrolyte properties of actin filaments might provide a deeper understanding of the fundamental mechanisms governing the intracellular ionic wave packet propagation in neurons. Infinitely long cylindrical models for actin filaments and approximate electrochemical theories for the electrolyte solutions were recently used to characterize these properties in *in vitro* and intracellular conditions. This article uses a molecular structure model for actin filaments to investigate the impact of roughness and finite size on the mean electrical potential, ionic density distributions, currents, and conductivities. We solved the electrochemical theories numerically without further approximations. Our findings bring new insights into the electrochemical interactions between a filament’s irregular surface charge density and the surrounding medium. The irregular shape of the filament structure model generated pockets, or hot spots, where the current density reached higher or lower magnitudes than those in neighboring areas throughout the filament surface. It also revealed the formation of a well-defined asymmetric electrical double layer with a thickness larger than that commonly used for symmetric models.

## 1. Introduction

Actin filaments and microtubules constitute a significant component of the cytoskeleton. Recently, they have gained a reputation as conducting nanobiowires forming an optimized information network capable of transmitting ionic waves in neurons [1, 2, 3, 4, 5, 6]. These polyelectrolytes are also potentially useful for the fabrication of dense arrays of electrical interconnections in 3D chips [7, 8], semiconductor nanowires using quantum dots - actin conjugation [9], bio-nanotransporters based on actin-gold nanowires 10, 11 and iontronic devices [12]. However, the underlying biophysical principles and molecular mechanisms that support the ionic conductance and transport along actin filaments are still poorly understood. Approximate theories using infinitely long cylindrical filament models have become a powerful tool to characterize the electrical conductivity properties of these polyelectrolytes. For instance, the high surface charge of the polyelectrolyte usually present in physiological conditions, causes an inhomogeneous arrangement of counter- and co-ions forming an electrical double layer (EDL) around its surface. The accumulation of these ions builds up an ionic conductivity and capacitance layer intrinsic to the EDL.[13, 14] Recently, we introduced a multi-scale approach for infinitely long cylindrical filaments in electrolyte solutions which has been shown to capture non-trivial contributions of the diffuse part of the EDL to the ionic conductivity and capacitance.[14, 15] These contributions revealed remarkable dependence on the electrolyte fluid composition, the filament electric surface potential, the ionic strength, and EDL width. As a result, the electrical conductivity in intracellular conditions showed lower values than those predicted for *in vitro* experiments, while the opposite result was obtained for the capacitance. Moreover, the effects of surface irregularities were neglected in these predictions. Additionally, the linear approximation to the Poisson-Boltzmann (PB) equation was used to calculate the EDL properties.

Certainly, more accurate approaches and realistic filament models might elucidate the global and the local ionic transport properties. In this work, we performed numerical calculations on a molecular structure for actin filaments using commercial software Comsol Multiphysics^®^ to investigate the impact of surface irregularities on the polyelectrolyte properties of these filaments in *in vitro* and intracellular conditions. We also implemented the non-linear Poisson-Boltzmann (NLPB) equation to more accurately account for the electrochemical effects on the surface of the filament and the EDL. For simplicity, we didn’t consider dissipative and damping forces. We determined the ionic conductivity, the capacitance, the radial and longitudinal current density profiles, the mean electric potential, the radial ionic concentrations, and the axial velocity profiles. Our results are compared with those obtained recently using an infinitely-long cylindrical model [14] to reveal the effects of the filament roughness and finite-size on these quantities.

## 2. Approach

The details on the hydrodynamic and electrical theories, and boundary conditions implemented in Comsol Multiphysics^®^ software are described in the appendix.

### 2.1. Molecular structure model for actin filaments

We retrieved a biological structure unit model and topology information for actin filaments from the RCSB Protein Data Bank in pdb format[16]. We used software Phenix[17] and Mathematica[18] to generate the molecular structure for the biological ensemble of 25 G-actin monomers. This structure was processed using the molecular visualization software VMD[19]. The STL triangular mesh was smoothed using the QuickSurface feature in VMD to increase computational efficiency. The mesh was subsequently imported into Comsol Multiphysics^®^[20]. (see Figure 1).The resulting filament geometry is characterized by a volume *V*_*f*_ = 5365700 Å^3^, a surface area *A*_*f*_ = 245700 Å^2^, and length *L*_*f*_ = 767.24 Å.

**Figure 1:**
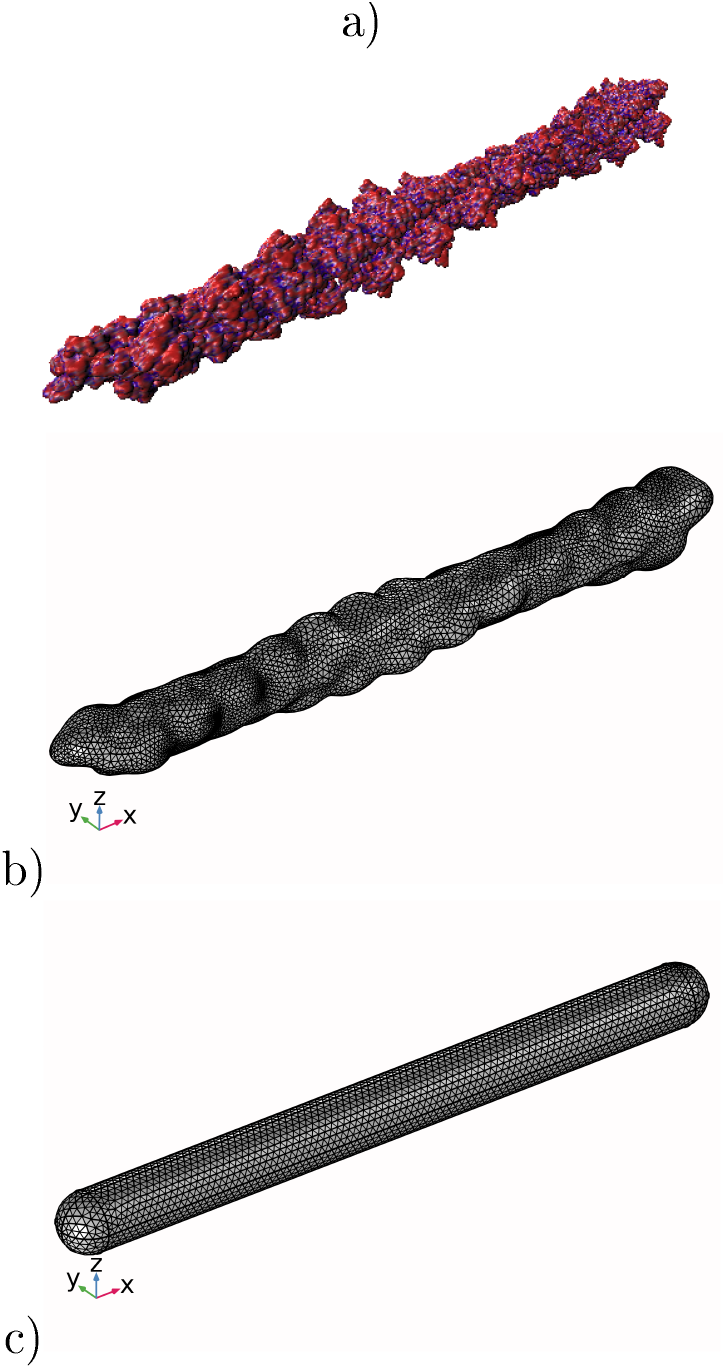
a) Molecular structure obtained using VMD1’s QuickSurf feature. b) Smoothened mesh imported into Comsol. c) Simplified (cylinder) model obtained using the dimensions of the smoothened mesh

Additionally, we used the default configuration and the Charmm force field[21] in the web application pdb2pqr[22] to add hydrogens and assign atomic charges and sizes into the filament molecular structure. The resulting output file in pqr format was used to sum all the atomic charges in the filament. This calculation yields a molecular structure charge of *Q*_*f*_ = −350*e, e* being the elementary unit charge. The filament surface charge was estimated assuming an homogeneous charge distribution as *σ*_*f*_ = *Q*_*f*_ */A*_*f*_ = −0.002282 *C/m*^2^.

### 2.2. Cylindrical filament and Electrolyte models

We also simulated in Comsol a spherocylindrical filament model with finite size *L*_*c*_ and effective radius *R*_*c*_ (see Figure 1c). A high geometrical and electrical similarity between the molecular structure and finite-size cylindrical models is achieved by setting *Q*_*c*_ = *Q*_*f*_ = −350*e, L*_*c*_ = *L*_*f*_ = 767.24 Å, and 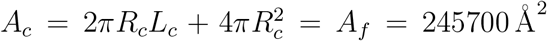. The latter relation yields to a value for the effective cylinder radius *R*_*c*_ = 45.557. Additionally, the finite size cylinder model preserves the molecular structure surface charge density, namely *σ*_*c*_ = *σ*_*f*_ = − 0.002282 *C/m*^2^.

We also consider the infinitely-long, azimuthally symmetric cylinder model introduced in reference [14]. In this work, we characterize the filament with the same surface charge density and radius characterizing the finite size cylinder for comparison purposes. Details about the governing equations and mathematical formulation of this model are described in reference [14].

The intracellular and *in vitro* conditions are defined in reference [[14]]. The molar concentration parameters for the intracellular solution are 140 mM K^+^, 4 mM Cl^−^, 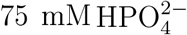, and 12 mM Na^+^ at 310, whereas for the *in vitro* solution are 100 mM K^+^ and 100 mM Cl^−^ at 298 K.

## 3. Results

The approach and parameters introduced in Section 2 were used in Comsol simulations for the molecular structure and finite-size cylindrical filament models to determine the radial and longitudinal current density profiles, the mean electric potential (MEP), the radial ionic concentrations, the axial velocity profiles, and the total current and resistance around the filament. We used software JACFC [23] to perform the corresponding numerical calculations for the infinitely-long, azimuthally symmetric cylinder model.

### 3.1 Mean electric potential

In Figure 4, we compare the MEP profiles at the cross-section plane location *x* = 0 (middle filament length) for the molecular structure and finite-size cylindrical filament models in intracellular electrolyte solutions. While the finite size model shows a high azimuthal symmetry on the MEP values, the molecular structure filament model displays a notorious angular dependence arising from the surface irregularities. Indeed, the filament’s shape results in larger or smaller radii depending on orientation. For instance, the shape of the filament at 240 ° results in a radius of approximately 58 Å, whereas the radius at 315 ° is almost 40% smaller (approximately 38 Å). This radius difference affects the MEP profile, as the smaller radius (38 Å at 315 °) yields an electronegativity of -22 mV on the surface, whereas the larger radius (58 Å at 240 °) reaches -19 mV. A similar asymmetry on the MEP values was also observed for *in vitro* conditions. A three-dimension MEP representation is given in Figures 9a and b for intracellular and in-vivo electrolyte solutions, respectively. The irregular shape of the molecular structure model generates hills and valleys where the MEP can reach higher or lower magnitudes than those in neighboring areas throughout the filament surface. Additionally, a colorful two-dimension representation of the asymmetric MEP behavior predicted by the molecular structure and finite-size cylindrical filament models is visualized in Figures 2 and 3 for intracellular and in-vivo conditions, respectively. These figures show the data obtained at the left border (*x* = *L*_*f*_ */*16), center (*x* = *L*_*f*_ */*2), and right border (*x* = 7*L*_*f*_ */*16) cross-sectional planes of the filament. These planes are defined as parametric surfaces described in Equation 13. With this setup, the MEP was calculated as a function of the polar angle, *θ*, and the radial separation distance *r*. Moreover, the yellow color represents the MEP value in bulk solution, whereas the blue color is the corresponding value inside and on the filament surface. In between, the color gradient represents the MEP values in the EDL. In the right columns, the white circles within the filaments come from the cylindrical to Cartesian coordinates transformation, as no MEP values were considered in these figures for distances shorter than 38Å. As a unique feature of this representation, the asymmetric behavior of the MEP is represented by the deviation of the contour line delimiting the different color layers from a vertical line (figures in column 1) or circle (figures in column 2) in polar or Cartesian coordinates, respectively. Clearly, the results for the finite size model (row 4) at the middle of the filament length show a high azimuthal symmetry on the MEP values. Whereas, the comparison between the figures in row 4 (finite size model) and the figures in rows 1-3 (molecular structure model) provides details about the impact of azimuthal and axial asymmetries of the molecular structure on the MEP values. Additionally, the comparison between each row in Figure 2 and the corresponding in Figure 3 reveals the effects of the electrolyte condition on the MEP values. Despite the filament irregular shape and electrolyte condition, Figures 2 and 3 reveal a well defined, asymmetric thin layer around the filament surface, similar in thickness to the one observed around the finite-size cylinder model. The color gradient width between the blue and yellow represents the EDL thickness, its value is (∼14Å) and remains approximately the same regardless of the axial location. Similar results are observed in *in vitro* conditions; however, the EDL thickness is slightly larger (∼18 Å) than the intracellular condition. This result is in agreement with the infinitely-long cylinder model predictions in which the exponential decay rate of the MEP in the EDL is proportional to the debay length.

**Figure 2:**
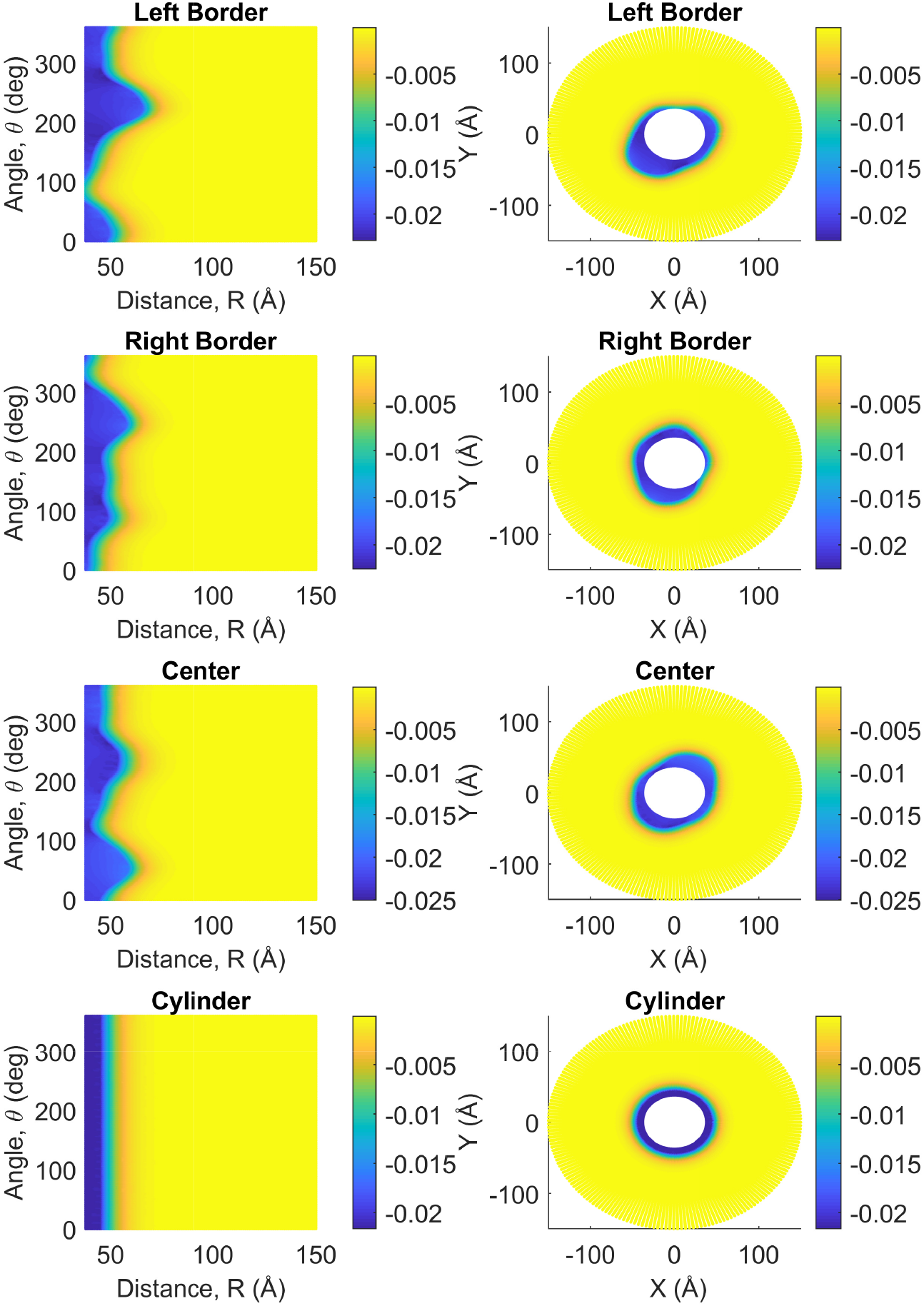
Measurements for intracellular electric potential (*V*) as a function of angle, *θ* (°), and separation distance, *R*(Å), on the left column and as cross-sectional views perpendicular to the filament’s longitudinal axis on the right column..Both columns include measurements for the filament at the center, left, and right borders, as well as at the center of the cylinder model.

**Figure 3:**
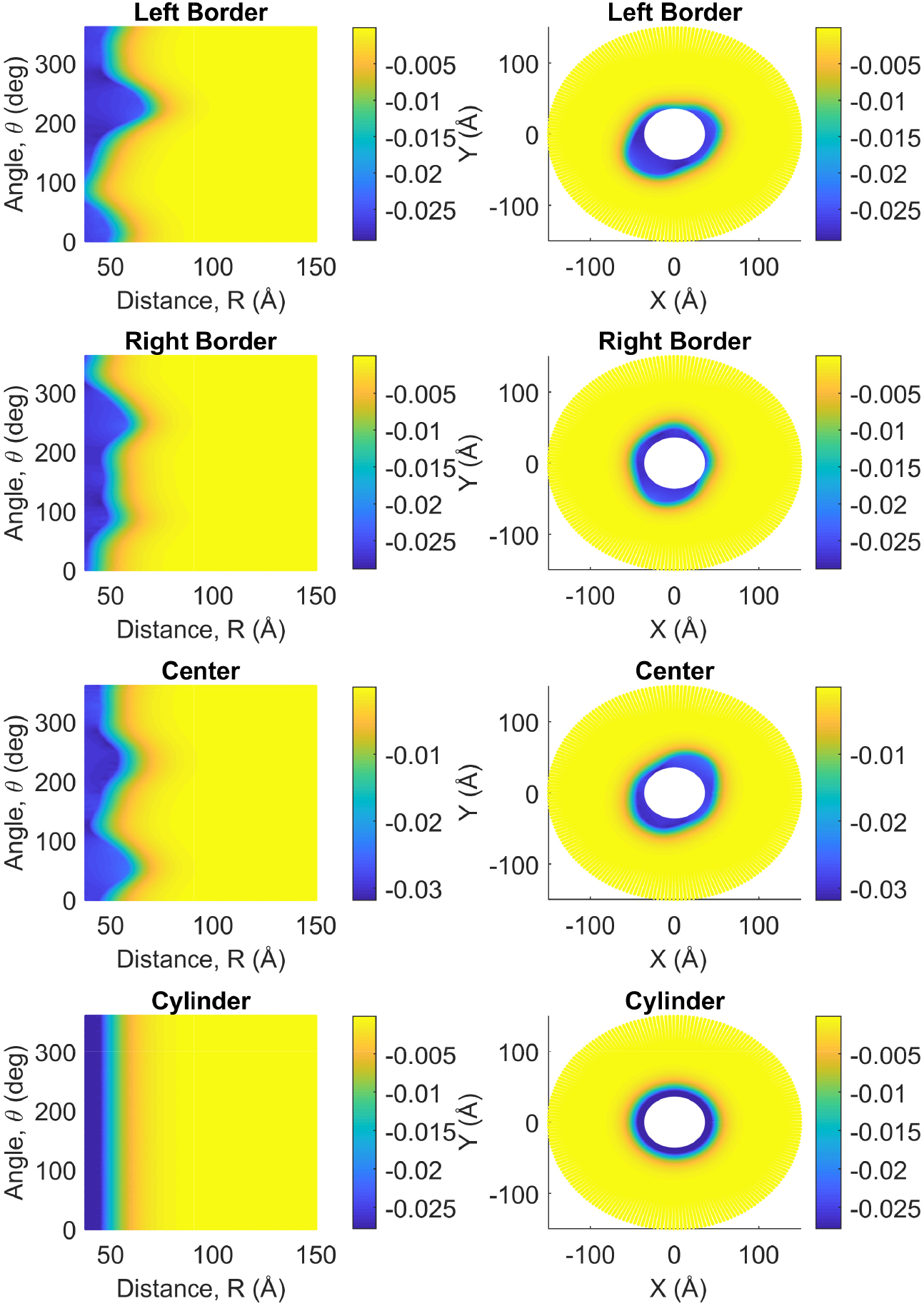
Measurements for *in-vitro* electric potential (*V*) as a function of angle, *θ*(°), and separation distance *R*(Å),on left column,and as cross-sectional views perpendicular to the filament’s longitudinal axis on the right column. Both columns include measurements for the filament at the center,left, and right borders,as well as at the center of the cylinder model.

**Figure 4:**
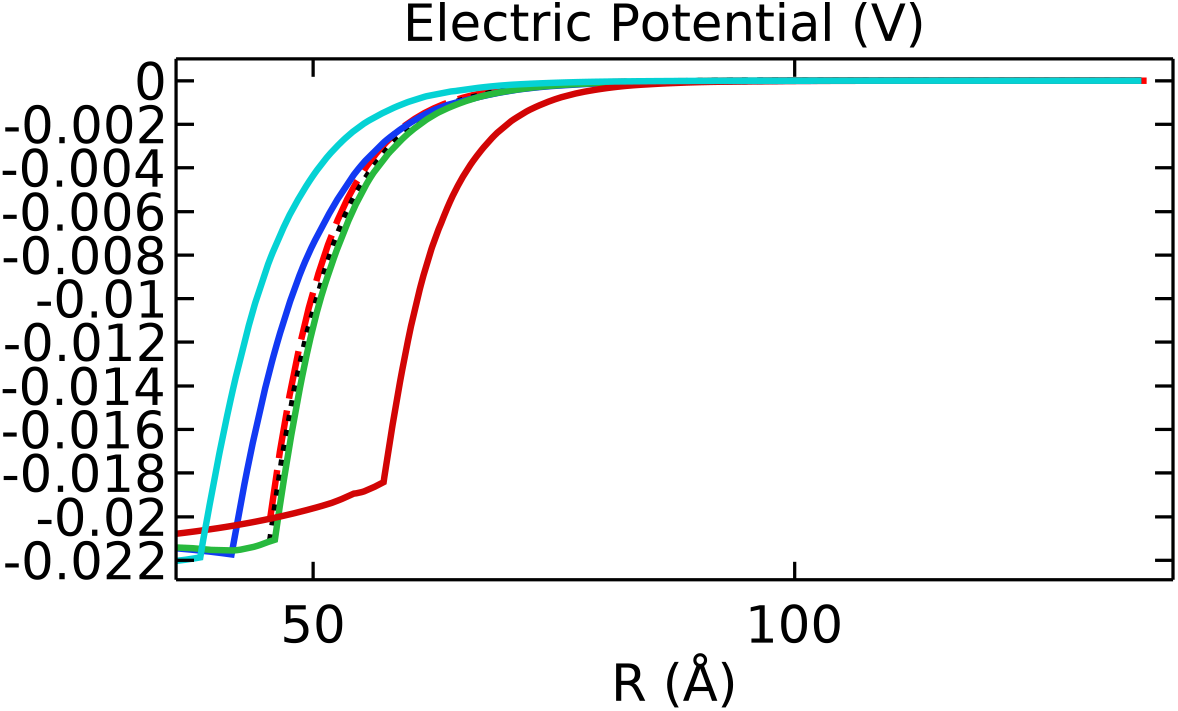
Mean Electric potential (*V*) as a function of the separation distance *R*(Å) obtained for the three models using intracellular electrolyte conditions: Analytical solution (magenta dashed line), simplified cylinder solution (black dash-dotted line) and more realistic filament solution (solid lines). Measurements for the filament were obtained at three different angles: 60° (blue), 135° (green), 240° (red), 315° (cyan).

Additionally, Figure 5 shows the comparison between the azimuthally and axially aver-aged MEP obtained for the molecular structure and finite-size cylindrical filament models, and the result for the infinitely-long, azimuthally symmetric cylinder model under intracellular and *in vitro* conditions.

**Figure 5:**
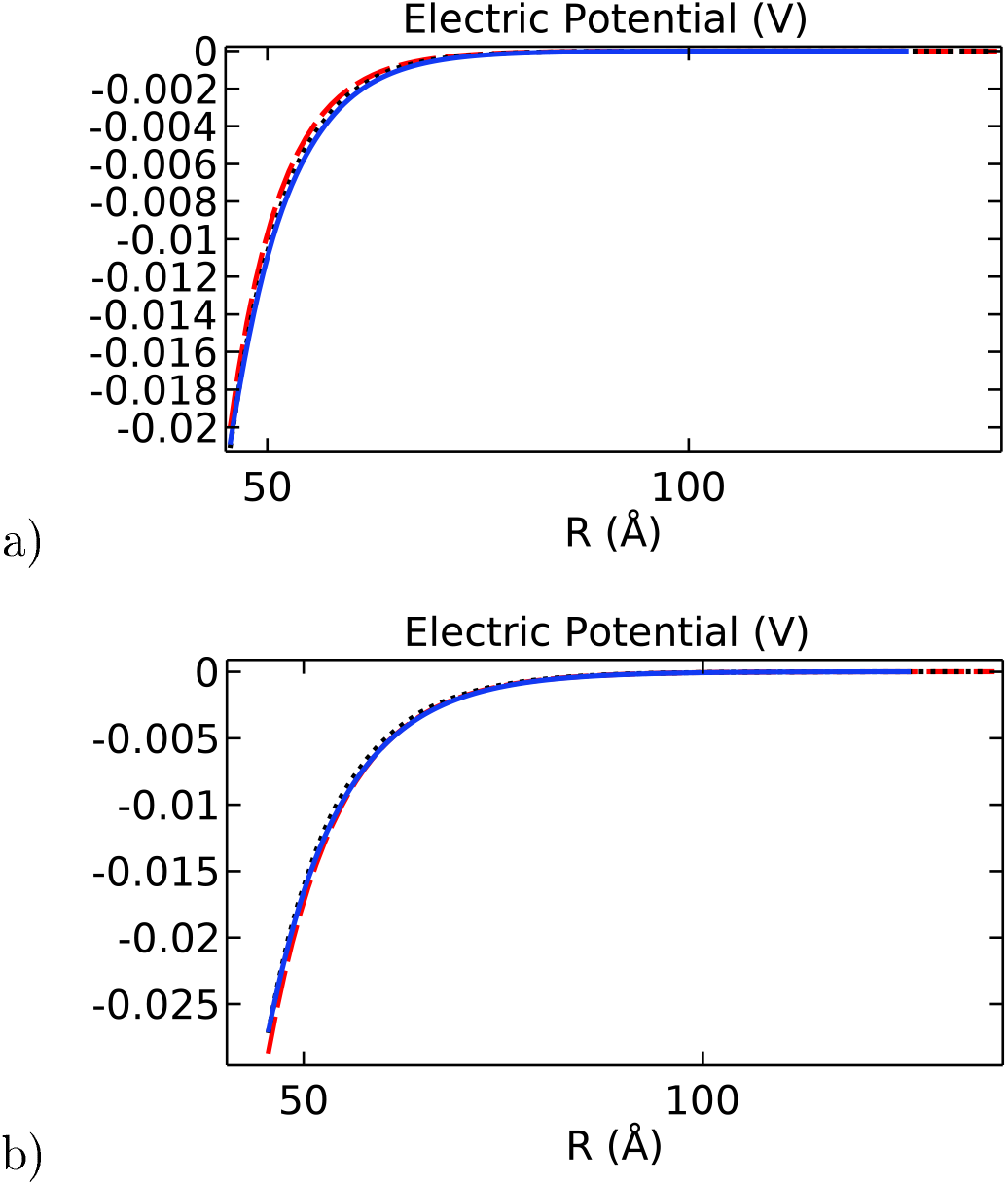
Mean Electric potential (*V*) as a function of the separation distance *R*(Å) for a) Intracellular and b) *in vitro* conditions. In both graphs, the blue solid line is the angular average obtained for the filament, the black dotted line is the solution obtained for the cylinder model, and the red dashed line is the analytical solution

It can be seen that the solution obtained for the infinitely-long cylinder model and that averaged for the finite-size cylinder models are in very good agreement. As a result, the filament length considered in this work (even larger) for the cylindrical model does not play a significant role in the Poisson-Boltzmann equation solution. While it reproduces the trend, the averaged molecular structure model solution for MEP in intracellular conditions is slightly higher in magnitude compared with those obtained for the other two models. Whereas, a lower magnitude is noticed in *in vitro* conditions. Moreover, the averaged molecular structure model solution in intracellular condition decays slower than its *in vitro* counterpart. Thus, while the filament’s irregular shape does have a local impact on the MEP (Figure 4), the averaged (global) electrical properties of actin filaments can be described accurately using azimuthal and axial symmetric cylindrical models.

### 3.2. Radial ionic concentration profiles

In comparison to the angular dependence observed on the MEP magnitude (figure 4), the impact of the filament’s irregular shape on the ion concentration profiles around the filament surface is broaden. This is because the inhomogeneous ionic concentration distributions in the EDL obey the Boltzmann statistics (Equation 11) where counter-ions accumulation and co-ions depletion around the filament surface depends exponentially on the MEP and the ionic valence magnitudes. An illustrative example of this phenomenon is given in Figure 6.a where potassium concentration profiles are plotted as a function of the radial separation distance from the filament axis. Each curve represents the concentration profile for a given orientation at the left border cross-section plane of the filament (*x* = *L*_*f*_ */*16). According to the asymmetric impact observed on the electric potential, there are higher number of ions accumulated in the regions where the distance between the axis and the surface of the filament is smaller and vice versa. For instance, the potassium concentration on the filament surface is ∼320 mM and ∼280mM at 60° and 240° respectively. A colorful representation of the asymmetric ion concentration behavior in polar and Cartesian coordinates is visualized in Figure 6.b. The asymmetric accumulation of potassium ions in the EDL is given by the deviation of the contour curve delimiting the yellow and blue zones from a vertical line or circle in polar or Cartesian coordinates, respectively. Overall, the asymmetric effects vanish in the bulk solution for radial distances larger than 70Å.

**Figure 6:**
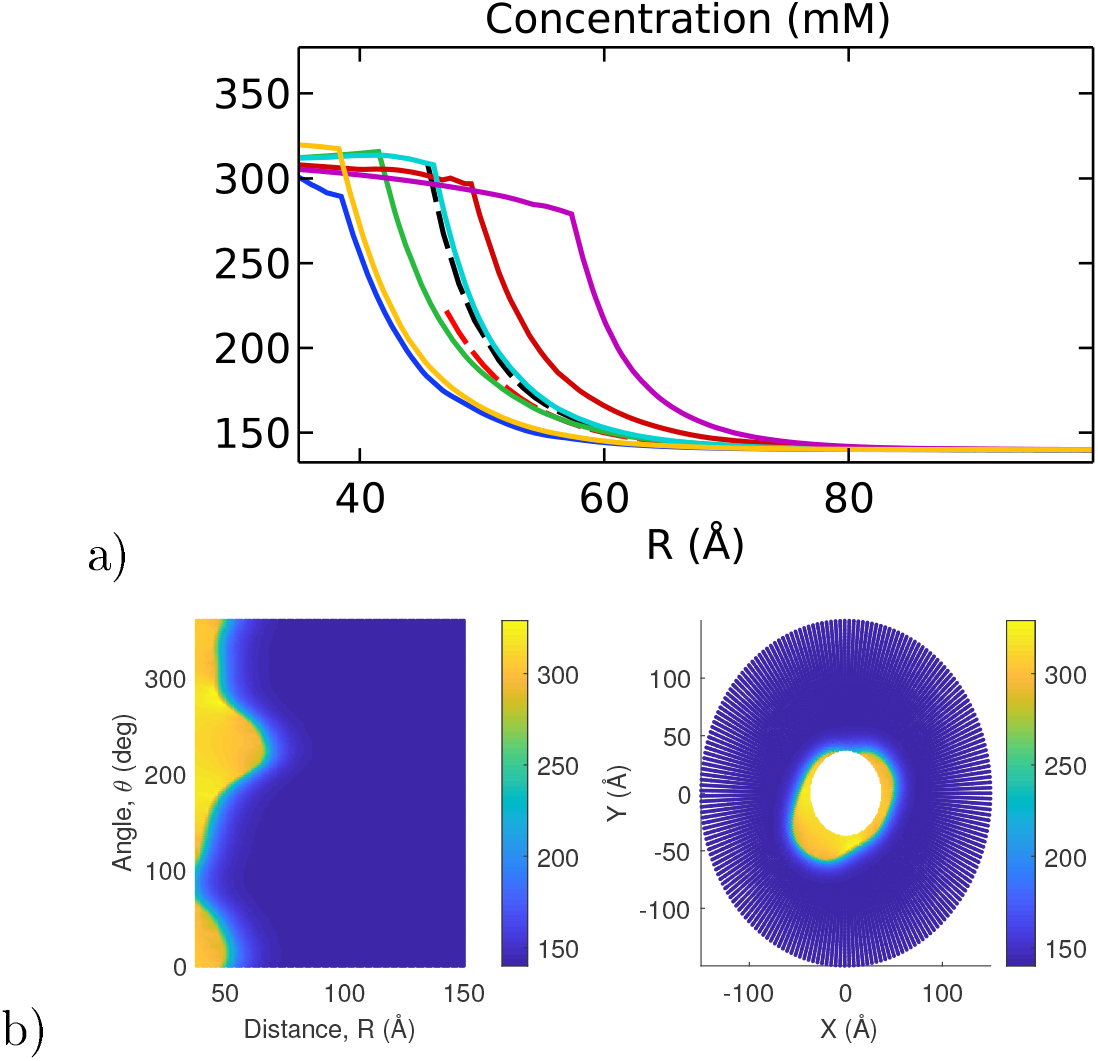
Angular comparison of potassium concentration on the surface of the filament at the left border of the filament (*x* = *L*_*f*_*/*16): a) Line graph showing the concentration profile for the filament at different angles (solid lines), as well as the profiles obtained from the cylindrical model (black dashed line) and the theoretical model (red dashed line). The evaluation angles are 0°(blue), 60° (green), 90° (red), 135° (cyan), 240° (magenta), 315° (yellow). b) Concentration around the filament surface as a function of distance and angle (right) and in Cartesian coordinates (left)

Similarly to the analysis done in Figure 5 on the MEP, Figure 7 shows the comparison between the azimuthally and axially averaged ionic concentration distributions obtained for the molecular structure and finite-size cylindrical filament models, and the result for the infinitely-long, azimuthally symmetric cylinder model under intracellular and *in vitro* conditions. In this case, the results obtained for the theoretical show ionic concentration levels at the surface that underestimate those values obtained for the molecular structure and finite-size cylindrical models. These differences are found in both intracellular and *in vitro* solutions, but are more significant for ion species with large concentration gradients (see, for instance, the curve for potassium ions in Figure 7.a). This discrepancy is partially due to the approximations introduced in the theoretical model. Additionally, small differences between the symmetric and asymmetric models prediction on the MEP values may generate substantial deviations for the ionic concentration distributions, since the asymmetric contributions generated in the molecular structure model from the MEP values are increased exponentially. As a result, these asymmetric contributions may be partially canceled out when averaging over the angular and axial coordinates.

**Figure 7:**
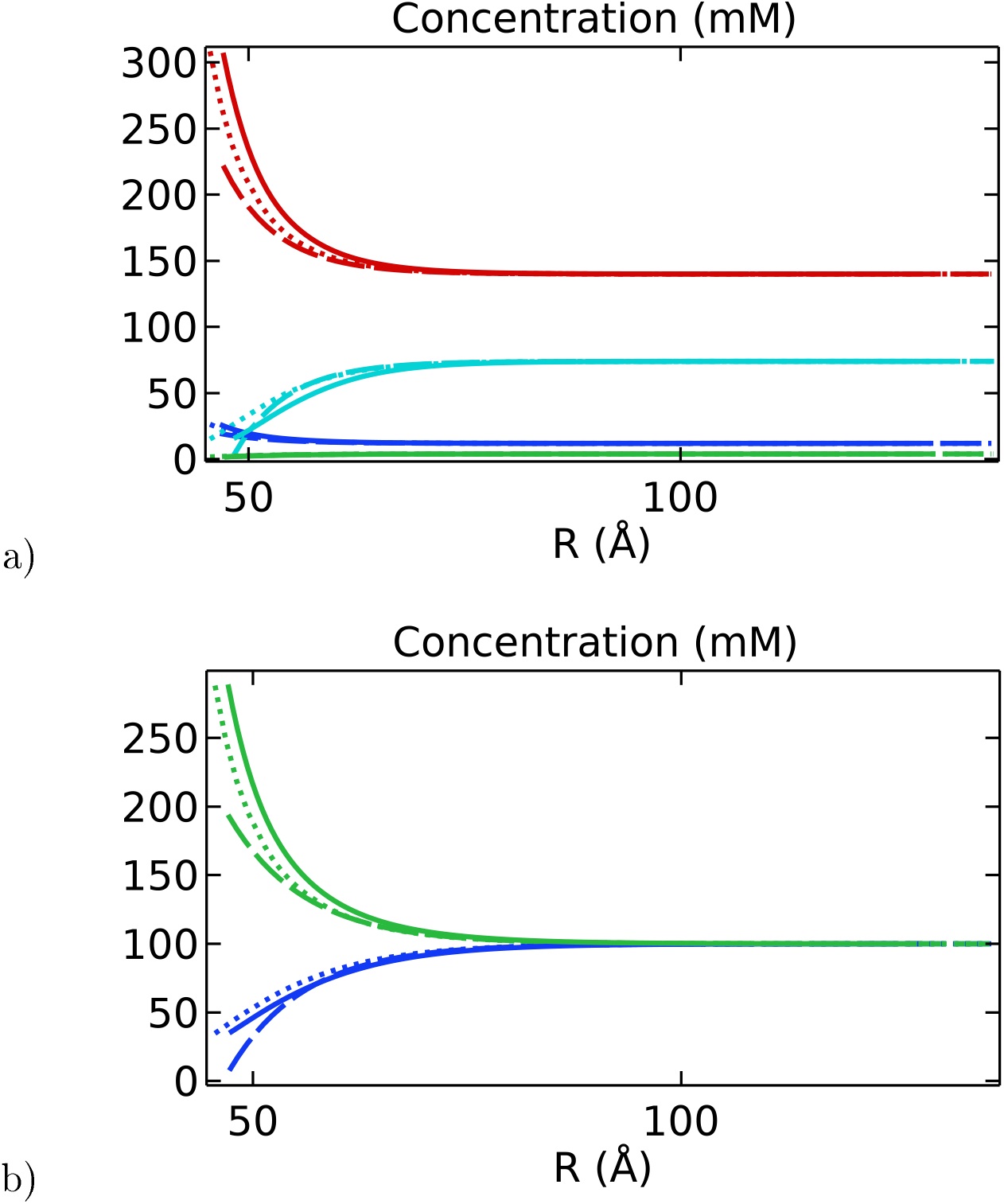
Ionic concentrations (*mM*) as a function of the separation distance *R*(Å) for a) intracellular and b) *in vitro* electrolyte solutions. In both figures, the solid line is the angular average obtained for the filament, the dotted line is for the cylinder, and the dashed line is for the analytical solution. Moreover, the ionic species are differentiated by color: Blue is sodium (Na), green is chlorine (Cl), cyan is hydrogen phosphate (HPO_4_), and red is potassium (K).

### 3.3. Axial velocity profiles

The velocity of the surrounding electrolyte is driven by the electrochemical interaction between the actin filament and the surrounding media. The electrolyte representing an intracellular environment has a higher ionic bulk density when compared with its *in vitro* counterpart, making it more difficult for the different ions to move and, therefore, reaching a lower bulk velocity than the one predicted for the *in vitro* solution.

Figure 8 displays the averaged velocity profiles predicted by the three models as a function of the radial separation distance. While the theoretical model predicts a bulk velocity of 0.43 m/s for the intracellular electrolyte, the molecular structure and cylinder models predict a bulk velocity closer to 0.46 m/s. This discrepancy is even smaller in the case of the *in vitro* electrolyte. This small difference can be attributed to the approximations used in each model. The analytic expression for the velocity profile in the theoretical model is proportional to the MEP, and the asymptotic value for large separation distances represents the uniform velocity magnitude in the bulk solution. This quantity depends on the zeta potential which is the MEP value at the slipping plane position. Since this position is not well defined in this model, it is usually approximated as the sum of the cylinder and the ion radii. While this approximation is not implemented in the other two models, the system is numerically solved in Comsol using a finite box size. As it was noted for the averaged MEP results, the molecular structure model further showcases a slower decay with the separation distance arising from the average effect of the surface irregularities. These differences between models, however, become negligible in the current density calculations, as discussed in Section 3.4.

**Figure 8:**
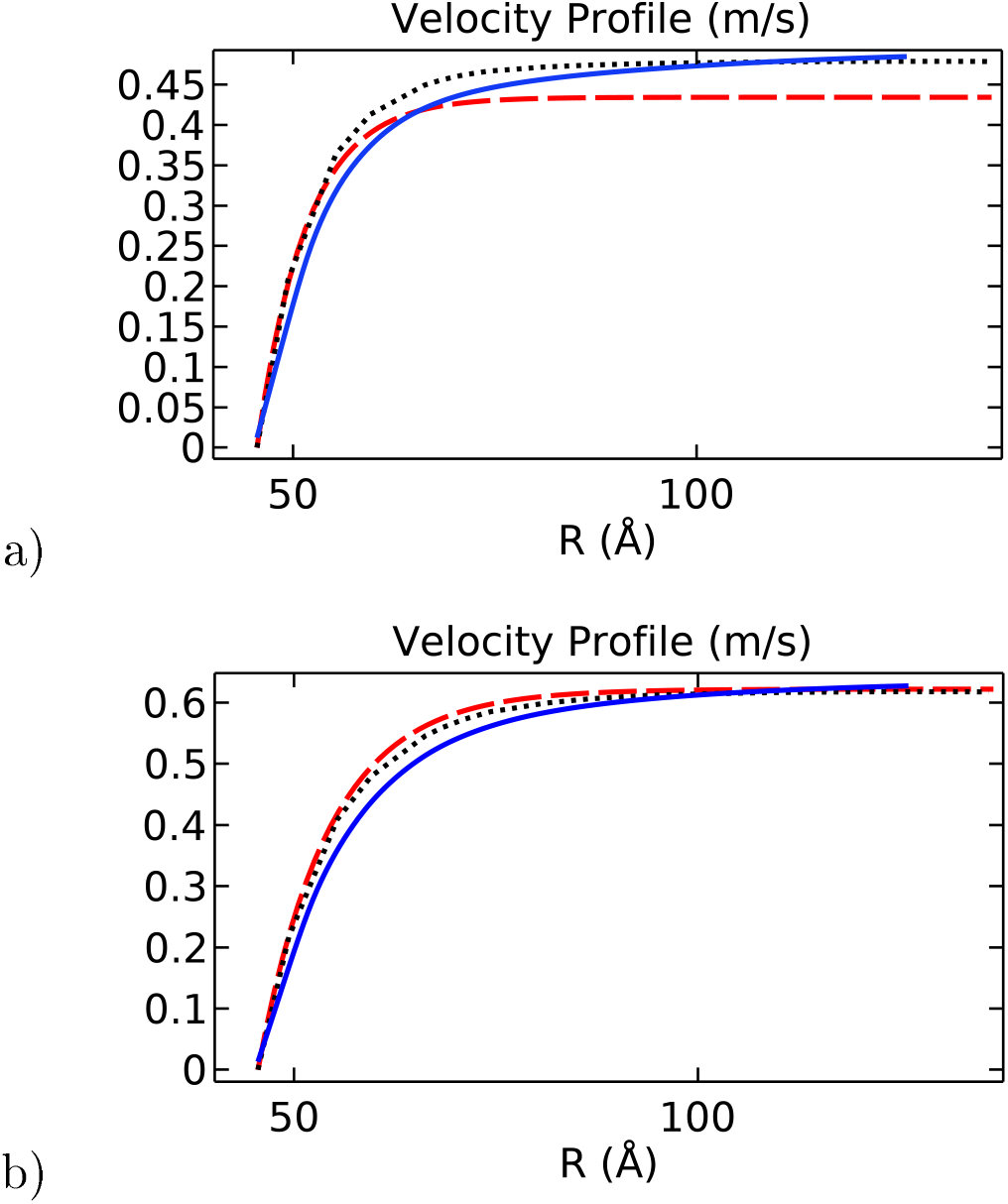
Velocity profile (*m/s*) as a function of the separation distance *R*(Å) for a) intracellular and b) *in vitro* electrolyte solutions. In both graphs, the blue solid line is the angular average obtained for the filament, the black dotted line is the solution obtained for the cylinder model, and the red dashed line is the analytical solution

### 3.4. Radial and longitudinal current density profiles

In this section, the solutions obtained for the MEP, and the velocity and ionic concentration profiles, are used to calculate the current density profiles (Equation 12).

Figure 9 shows a three-dimension illustration on the MEP and ionic current density around the filament molecular structure. The arrow field represents the total current density’s direction and magnitude in the EDL in intracellular and *in vitro* conditions. The roughness displays pockets; or hot spots; where the current density can reach higher or lower magnitudes than those in neighboring areas throughout the filament surface. It is also shown that the total current density vector changes the direction following the surface irregularities. These results indicate that the filament molecular structure plays a fundamental role in the local distribution and transport of ions along the surface.

**Figure 9:**
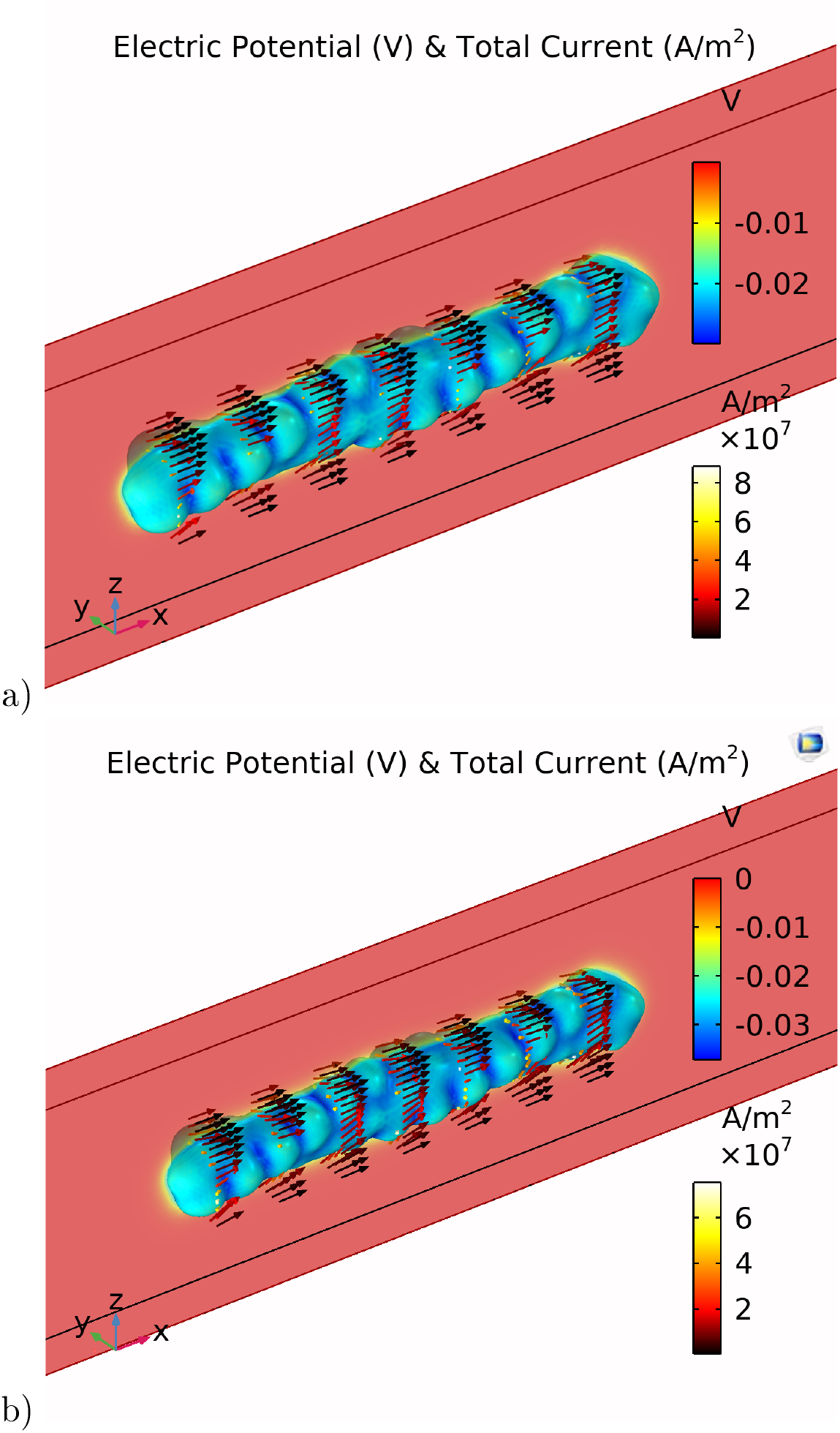
Total current density (*A/m*^2^) and electric potential (*V*) around the filament model for a) intracellular and b) *in vitro* conditions. In both figures, the arrow field indicates the direction and magnitude of the total current density (heat map colors) and the surface color indicates the electric potential (rainbow colors)

We analyzed the ionic current density vector predicted by the molecular structure and finite-size cylindrical models in terms of its longitudinal and radial direction components to facilitate the comparison with those theoretical model predictions. With the two-dimension colorful representation in polar and Cartesian coordinates used for the MEP, Figures 10 and 11 illustrate the asymmetric longitudinal current density behavior for intracellular and *in vitro* conditions, respectively. In this case, the white contour line appearing in the figures represents the filament surface since there is no current inside the filament’s domain. Alike the EDL displayed in Figures 2 and 3, the color gradient shows an asymmetric thin layer formed in the current density components around the filament surface. This is because the current densities depend primordially on the MEP values. Additionally, the behavior of the longitudinal current density varies drastically depending on the electrolyte type. Analogous to the MEP magnitude behavior, the longitudinal current density values decrease monotonically from the filament surface in *in vitro* conditions. However, it drops significantly within the first 10Å away from the surface, followed by a continuous increase until reaching the bulk value in intracellular conditions. This behavior may be in part due to the balance and competition arising in intracellular conditions between the electro-osmotic and convection contributions given in Equation 8 to calculate the longitudinal current density. These discrepancies are clearly manifest in Figures 12 and 13, which show the azimuthal and axial average of these components and compare them with those results obtained for both the infinitely-long and finite-size, azimuthally symmetric cylinder models.

**Figure 10:**
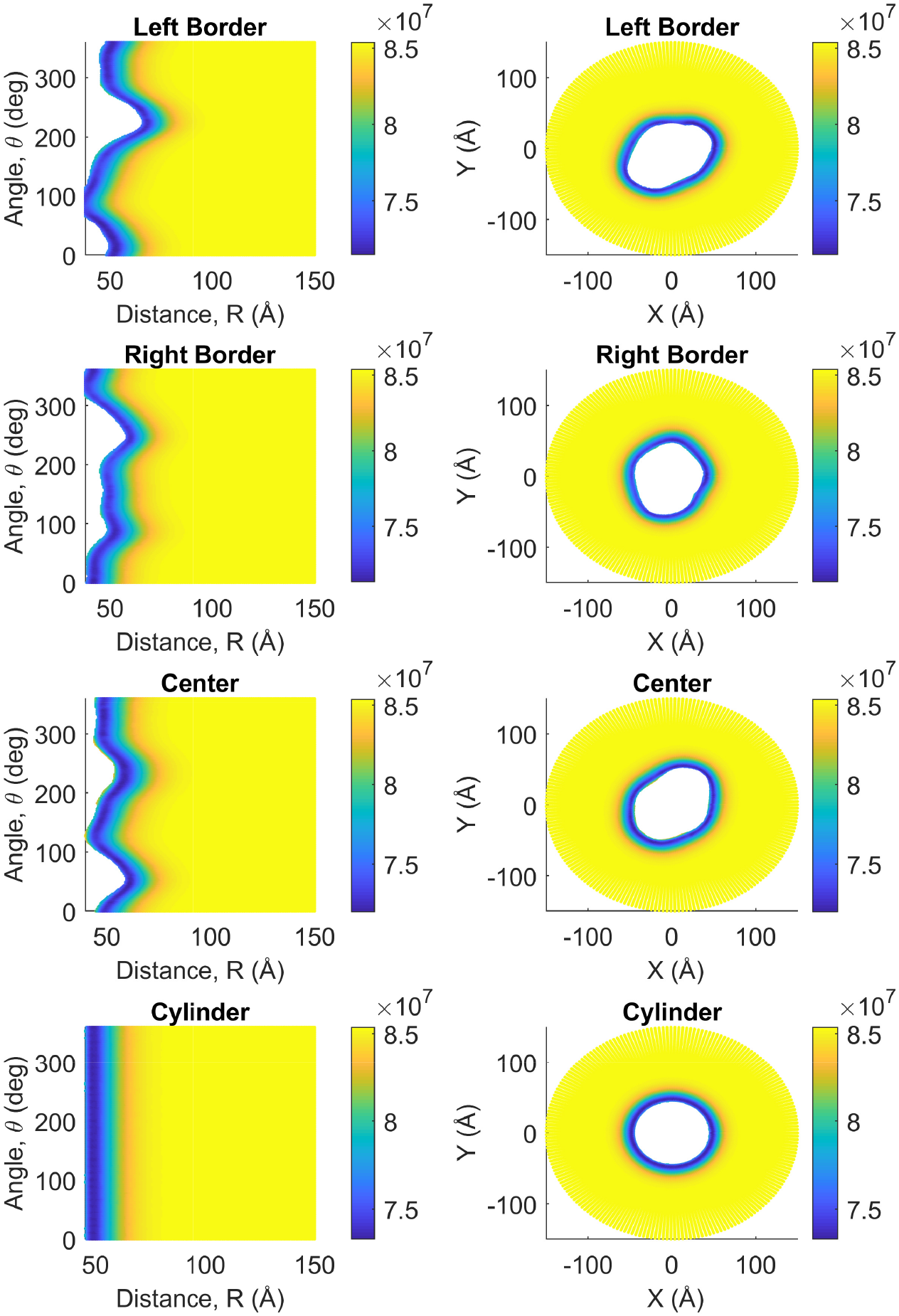
Intracellular longitudinal current density (*A/m*^2^) as a function of angle, *θ*(^°^), and separation distance, *R*(Å), on the left column, and as cross-sectional views perpendicular to the filament’s longitudinal axis on the right column. Both columns include measurements for the filament at the center, left, and right borders, as well as at the center of the cylinder model.

**Figure 11:**
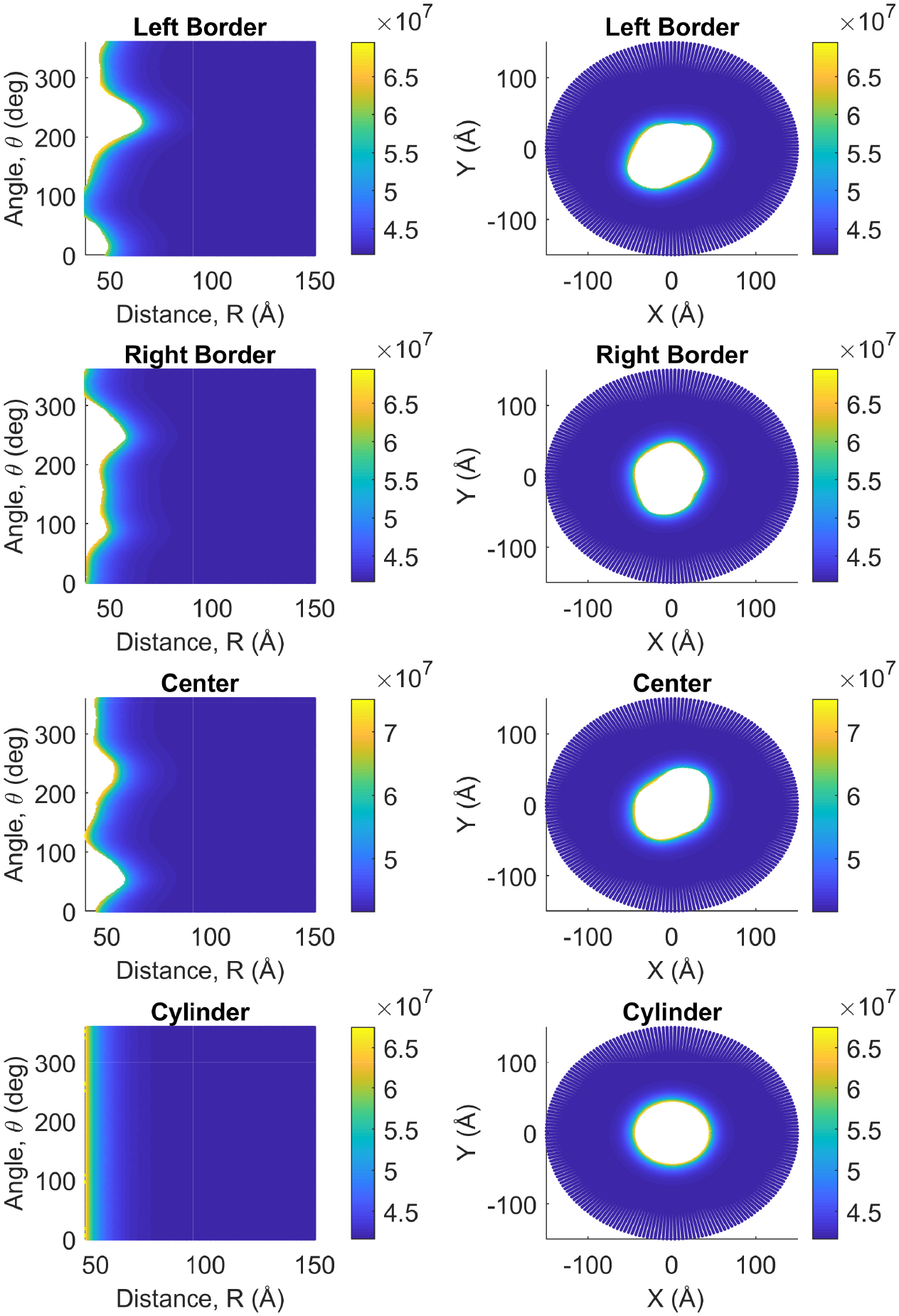
*In-vitro* longitudinal current density (*A/m*^2^) as a function of angle *θ*(°), and separation distance *R*(Å), on the left column,and as cross-sectional views perpendicular to the filament’s longitudinal axis on the right columns include data for the filament at the center left boders,as well as at the center of the cylinder model. The white space with in the filament and cylinder (right column) is due to the transformation from cylindrical to Cartesian coordinates, as no data were taken inside the filament is domain.

**Figure 12:**
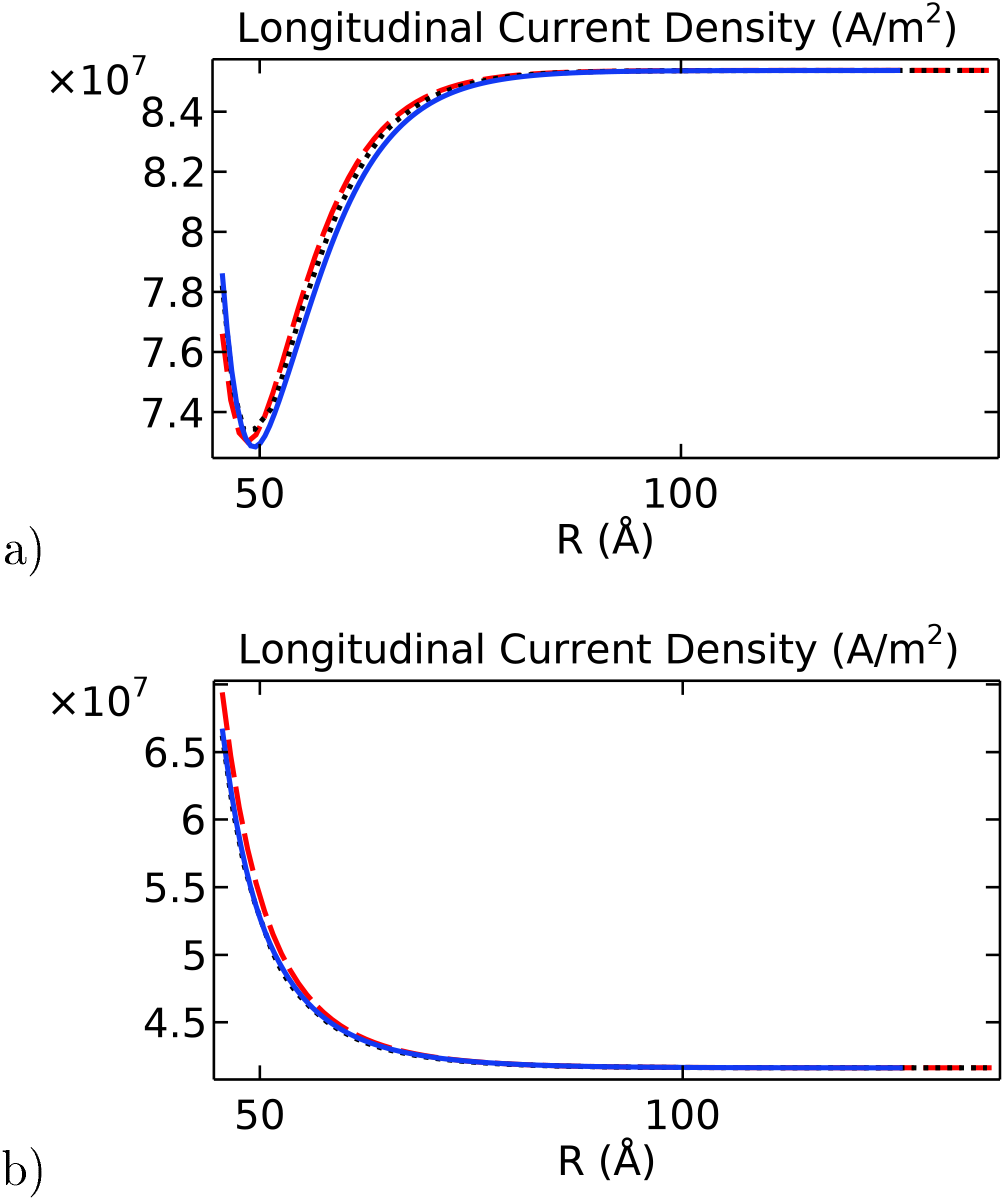
Longitudinal current density (*A/m*^2^) as a function of the separation distance *R*(Å) for a) intracellular and b) *in vitro* electrolyte solutions. In both graphs, the blue solid line is the angular average obtained for the filament, the black dotted line is the solution obtained for the cylinder model, and the red dashed line is the analytical solution.

**Figure 13:**
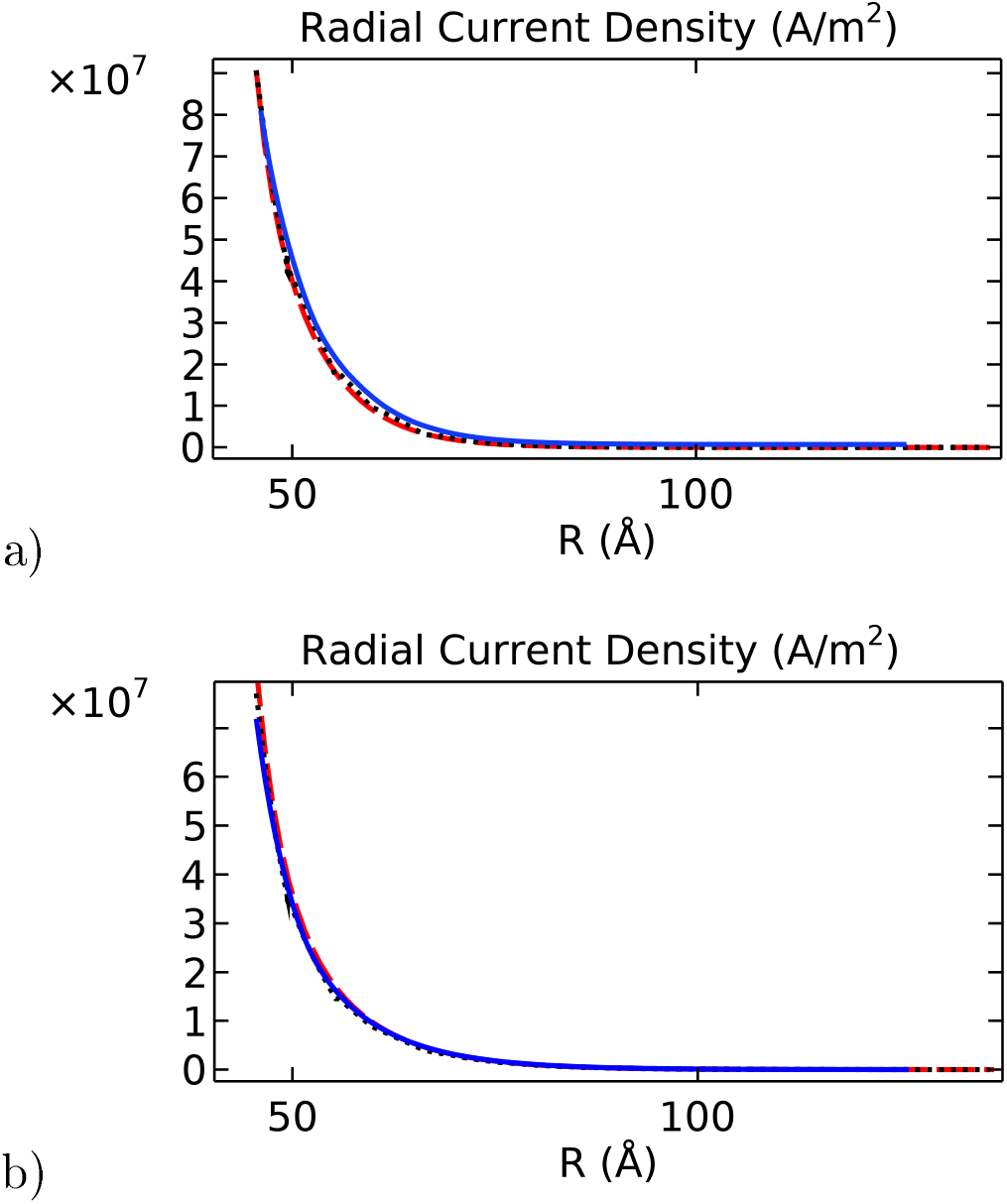
Radial current density (*A/m*^2^) as a function of the separation distance *R*(Å) for a) intracellular and b) *in vitro* electrolyte solutions.. In both graphs, the blue solid line is the angular average obtained for the filament, the black dotted line is the solution obtained for the cylinder model, and the red dashed line is the analytical solution

It is noted that despite the differences pointed out earlier about velocity, electric potential, and concentration profiles, the results obtained with the finite-size cylinder model and the theoretical model are in close agreement for both longitudinal and radical current density profiles under both intracellular and *in vitro* solutions. On the other hand, the molecular structure model results differ from their cylindrical counterparts on the *in vitro* radical current density profile,mainly (Figure 13.b). Since the radical current density depends basically on the MEP gradient values (Eq.8),these differences arise from similar deviations displayed in Figure 3 for the averaged MEP in *in vitro* conditions.

### 3.5. Total ionic currents and resistances

The radial and longitudinal current density solutions for the finite-size cylinder and molecular structure models were integrated over the polar angle and the radial distance to obtain the total ionic current values at each of the fifteen cross-sectional equispaced planes (Equation 16). Subsequently, we used Ohm’s law to calculate the total resistances. In the longitudinal direction, the voltage drop is given by the external voltage stimulus imposed on the system (Δ*V* = 0.15 V). In contrast it is calculated in the radial direction as the electric potential difference between the filament surface and the outer plane of the EDL. The results for the resistance along the molecular structure axis are shown in Figure 14. While the radial and longitudinal resistances behave uniform under intracellular electrolytes the results in *in vitro* show increased resistances at both ends of the filament.

**Figure 14:**
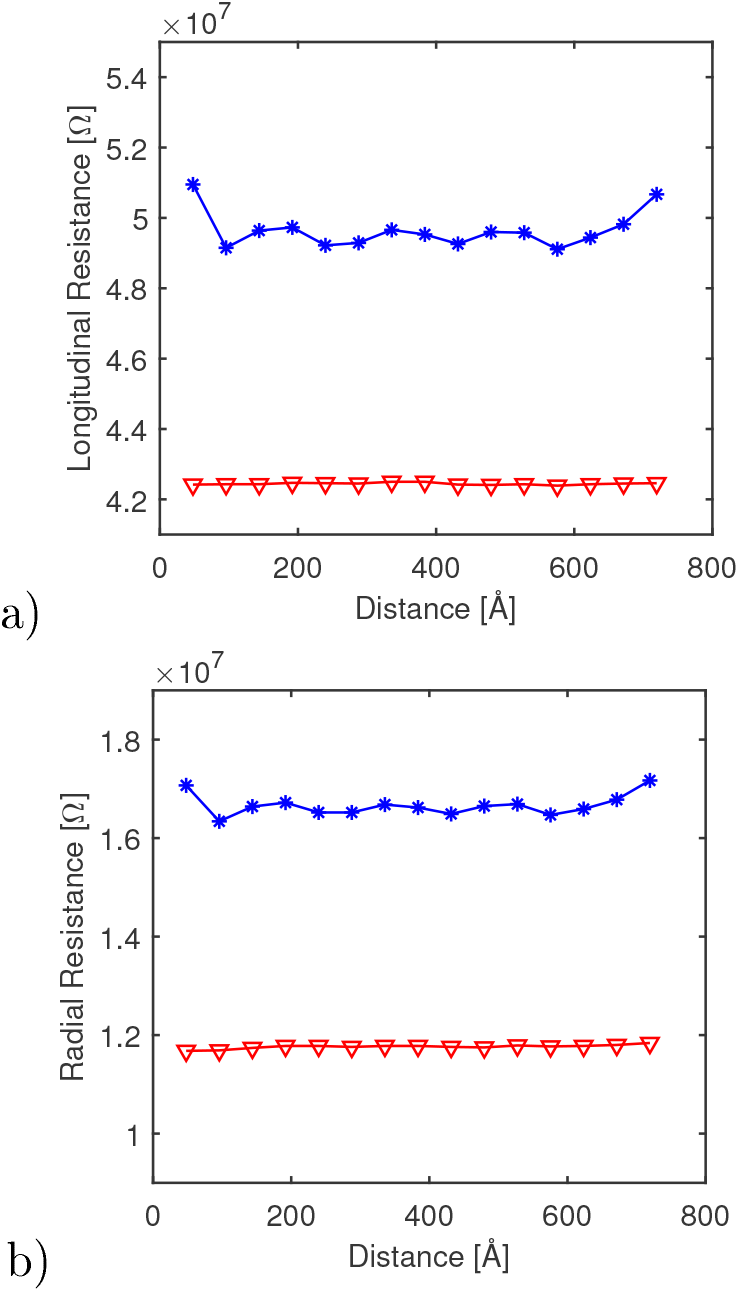
Resistance values along the filament axis in the a) longitudinal and b) radial directions. In both graphs, the blue line indicates *in vitro* conditions and the red line indicates intracellular conditions.

This resistance increase represents, at most, a 3% deviation from the overall average resistance throughout the filament axis. This small difference is found in both radial and longitudinal directions. Furthermore, the effect of the ends in *in vitro* electrolytes is not replicated when investigating the results of the finite-size cylinder model. Therefore, we attribute these discrepancies to the reduced electrochemical interaction occurring at the filament ends, leading to a lower current density and, thus, a higher resistance in that area (see left and right border panels of Figure 11). Due to the slight deviation incurred by this increase, we consider the effects of the filament ends negligible in the total current density and resistance calculations.

The total current density and resistance for the molecular structure and finite-size cylinder models were calculated by averaging their values over the fifteen cross-sectional equispaced planes. We used software JACFC [23] to calculate these quantities for the infinitely-long, azimuthally symmetric cylinder model. The results for the *in vitro* and intracellular conditions are reported in Tables 1 and 2, respectively.

**Table 1:** Total current and resistance comparison between filament, cylinder, and theoretical models for the *in vitro* case

| Model | Longitudinal<br>Current<br>[$10^{-9}$A] | Longitudinal<br>Resistance<br>[$10^7\Omega$] | Radial<br>Current<br>[$10^{-9}$A] | Radial<br>Resistance<br>[$10^7\Omega$] |
| --- | --- | --- | --- | --- |
| Filament | 3.02 | 4.96 | 1.40 | 1.67 |
| Cylinder | 2.99 | 5.01 | 1.37 | 1.66 |
| Theory | 3.06 | 4.91 | 1.49 | 1.68 |

**Table 2:** Total current and resistance comparison between filament, cylinder, and theoretical models for the intracellular case.

| Model | Longitudinal<br>Current<br>[$10^{-9}$A] | Longitudinal<br>Resistance<br>[$10^7\Omega$] | Radial<br>Current<br>[$10^{-9}$A] | Radial<br>Resistance<br>[$10^7\Omega$] |
| --- | --- | --- | --- | --- |
| Filament | 3.53 | 4.24 | 1.56 | 1.18 |
| Cylinder | 3.56 | 4.21 | 1.47 | 1.21 |
| Theory | 3.46 | 4.33 | 1.48 | 1.21 |

Overall, the results obtained in intracellular conditions for the finite-size cylinder and the molecular structure models are in very good agreement with those obtained for the theoretical model. With the most significant deviation being 5% in the total radial current obtained for the molecular structure model, we concluded that the shape impact on the global electrochemical phenomena occurring at and around the filament surface is minimal. Similarly, the total radial current obtained for the finite-size cylinder model in conditions resulted in 8% deviation from the theoretical model prediction. This deviation, nonetheless, was compensated by the MEP as the total radial resistance with both electrolytes did not surpass 3%. Additionally, the total longitudinal currents and resistances obtained for the finite-size cylinder and molecular structure models with both electrolyte types did not exceed 3% deviation from the theoretical model results.

This comparison further establishes that a more realistic model provides a higher level of insight into the local behavior of the electrochemical phenomena occurring at and around the filament surface. Moreover, the cylindrical models can accurately describe the global trends for such electrochemical phenomena.

## 4. Conclusion

In this article, we used a molecular structure model for actin filaments and electrochemical theories to investigate the impact of roughness and finite-size on their polyelectrolyte properties.

The approach was shown to capture non-trivial contributions to the mean electric potential and electrical conductivity arising from the interactions between an actin filament molecular structure and the surrounding ions accumulated around its surface. In particular, it revealed pockets, or hot spots, where the ionic current density and electric potential can reach higher or lower magnitudes than those in neighboring areas throughout the filament surface. It also predicted the formation of a well-defined asymmetric electrical double layer around the filament surface, similar in thickness to the one observed around the finite-size cylinder model. The EDL thickness predicted from the electric potential calculations for intracellular conditions is smaller than its *in vitro* counterpart. This finding agrees with the theoretical model predictions in which the MEP decay throughout the EDL is a function of the debay length.

The asymmetric contributions on the polyelectrolyte properties of actin filaments mostly canceled out when averaging over the angular and axial coordinates, which is a clear indication that utilizing a filament structure geometry is needed for understanding the elecro-chemical phenomena occurring on the molecular structure surface, while a simplified model provides sufficient information to determine the global behavior of the phenomena around the structure. Future directions of this work include the implementation of a time-dependent formulation to account for dissipative and dumping forces including explicit ionic particle sizes and solvent friction. We will also consider inhomogeneous filament surface charge densities to model the impact of the atomic charge distribution associated with the amino acid sequence throughout the molecular structure surface.

## 5. Confilcts of Interest

There are no conflicts of interest to declare

## 6. Acknowledgements

This work was supported by NIH Grant 1SC1GM127187-04.

## 7. Appendix

### 7.1. Hydrodynamic and electrical theories

The Electrostatic and Creeping Flow modules for equilibrium conditions were used to carry out the simulations for the molecular structure filament and the finite-length cylindrical models in Comsol Multiphysics^®^

The Electrostatics interface solves Gauss’ law for the electric field **E** and mean electric potential *V*

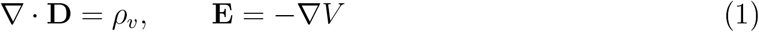

where **D** is the displacement field, and *ρ*_*v*_ = *F Σ c*_*i*_*z*_*i*_ [*C/m*^3^] is the space charge density defined over the electrolyte domain (*F* being the Faraday constant, and *c*_*i*_ and *z*_*i*_ being the ionic species molar concentration and charge, respectively). The displacement and electric fields are related by the constitutive relation **D** = *ϵ* _0_*ϵ* _*r*_**E**, where *ϵ*_*r*_ is the material relative permittivity (2 for the filament and 78.3 for the electrolyte solution). Furthermore, a constant surface charge density condition *ρ*_*s*_ = *σ* was applied to the filament surface, and a zero charge condition **n · D** = 0 was applied to the outer surface of the surrounding medium, due to the electroneutrality condition of the bulk solution.

The creeping flow interface solves Navier-Stokes equations for an incompressible flow. Given the nature of our system, we can neglect the inertial term and the pressure gradient, and assume an incompressible fluid. Thus making the governing equations be as follows

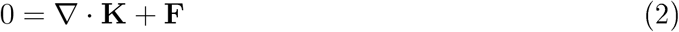

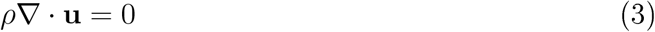

The first term on the right-hand side of Equation 2 refers to the viscous stress **K** = *η* ∇ · **u**, *η* is the fluid viscosity, **F** is the external electric force applied to the system per unit of volume, *ρ* is the fluid density, and **u** is the fluid velocity. In our system, the external electric potential differential (Δ*V* = 0.15*V*) drives the fluid motion along the filament axis; therefore, a uniform volume electric force condition is placed on the surrounding medium domain

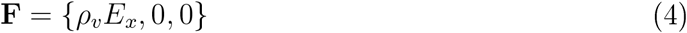

where *ρ*_*v*_ is the space charge density, and *E*_*x*_ = ∇*V/L*_*g*_ stands for the external electric field magnitude per unit of a monomer length (*L*_*g*_ = 54 [Å]). With no other force acting on the system, a free flow boundary condition was placed on the faces of the surrounding medium perpendicular to the filament

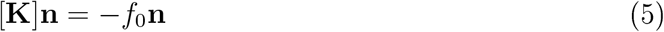

where the normal stress is set equal to *f*_0_ = 0 as the fluid is free to move in either direction. Additionally, two separate wall conditions were applied: a no-slip condition on the filament surface (Equation 6), and an electro-osmotic condition on the surfaces of the surrounding medium parallel to the filament (Equation 7),

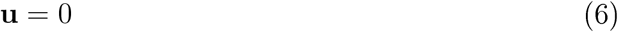

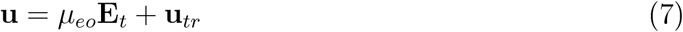

Here, 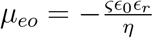 is the electro-osmotic mobility *(ς* is the zeta potential of the filament, defined as the average electric potential on the filament surface), and **E**_*t*_ = **E** − (**E · n**)**n** with **E** being the electric field solved for with the electrostatics formulation. Additionally, the fluid velocity far away from the filament is estimated by **u**_*tr*_ = *μ*_*eo*_*E*_*x*_.

### 7.2. Ionic current density characterization

The mean electric potential *V* and the axial velocity *v*_*x*_ profiles are used in Comsol to determine the radial and longitudinal current density profiles, the radial ionic concentrations, and the total current around the filament’s molecular structure and finitely-long cylinder models.

Considering the longitudinal axis of the computational filament model is along the X-axis, the current density in Cartesian coordinates is calculated as

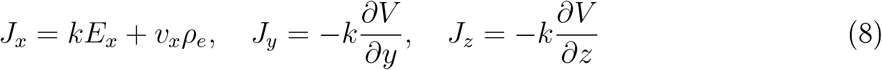

with

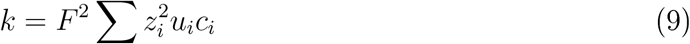

representing the electrolyte conductivity, and

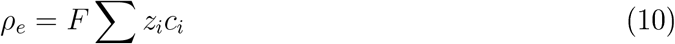

describing the total charge density distribution. Here, the parameter *F* represents the Faraday constant, and the parameters *z*_*i*_, and *u*_*i*_, stand for the valence and mobility of ionic species *i*, respectively. Additionally, the ionic concentrations are characterized by the Boltzmann distribution

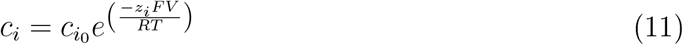

where 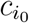 is the bulk concentration of species *i*, charge of species *i, R* is the gas constant, and *T* is the electrolyte temperature.

It is convenient to express the current density in polar coordinates to compare the results with the cylinder models. Using Equation 8, we can write

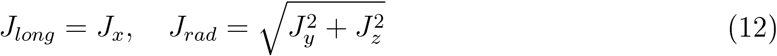

with a parametric surface defined as

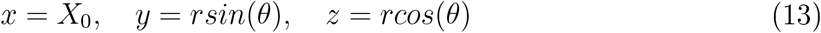

where *X*_0_ is a fixed position along the X-axis, *r* is any distance between the surface of the filament and 100 Å away from the surface, and *θ* is the angle measured around the filament surface (perpendicular to the filament axis) ranging between 0 and 2*π*.

### 7.3. Average and Integrals

To properly compare the theoretical, finite-size cylinder, and filament models, we implemented a coordinate system transformation in the filament model such that the surface of the filament would be located at the same distance as the radius of the theoretical and cylinder models. This transformation was implemented using Matlab as a shift in the raw data obtained from Comsol. Additionally, the Comsol measurements were obtained at fifteen different planes. These planes were perpendicular to the filament axis and equally-spaced throughout the filament. An illustration with three planes can be observed in Figure 15.

**Figure 15:**
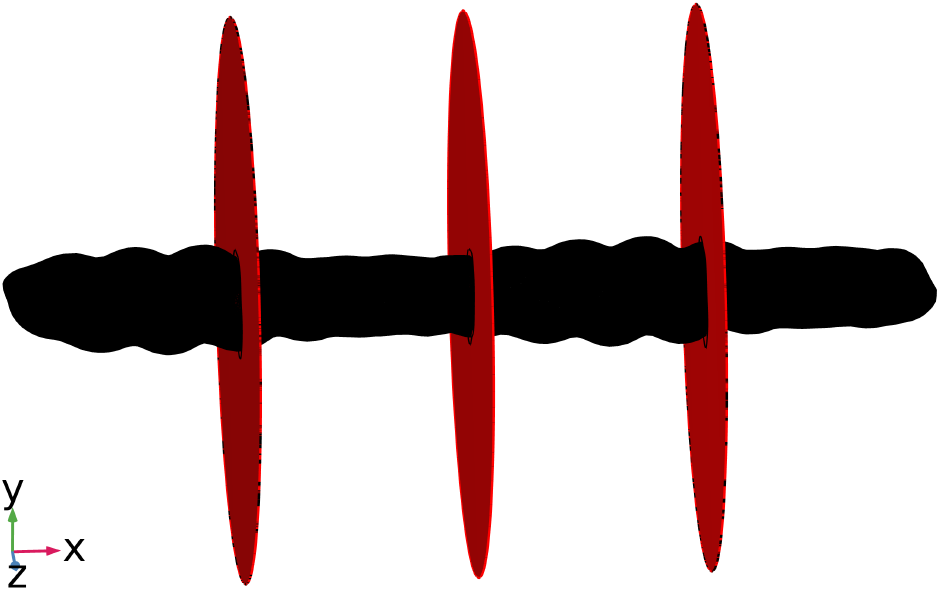
Three equally-spaced, perpendicular evaluation planes along the filament axis used to obtain electric potential, concentration, velocity, and current density values to calculate angular averages

As an illustrative example, we will describe these calculations for the intracellular electric potential (see Figure 2). At a given distance R near the filament surface, the electric potential will vary drastically as a function of the angle as observed on the left column of Figure 2. However, any measurement taken along the filament surface contour, will have a similar magnitude regardless of the angle, as observed in the right column of Figure 2. For this reason, the coordinate system transformation enabled the angular average calculation which, in the case of the electric potential, is determined as

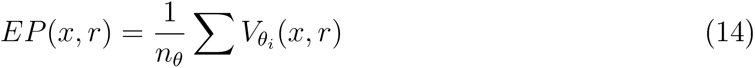

Where *EP* (*x, r*) is the electric potential as a function of distance along and perpendicular to the filament axis, *n*_*θ*_ = 200 is the number of angles used in the measurement, and 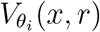 is the electric potential measured following the filament surface contour. This calculation was performed at each one of the fifteen planes (determined by *x*) and for contour surfaces at different distances away from the filament surface (determined by *r*) to obtain the average quantities discussed in Section 3. Moreover, given the filament model asymmetry, we calculated the average between all fifteen planes to obtain the overall average of the electric potential throughout the filament model. This overall average is used to compare the three models in the following sections and is expressed as.

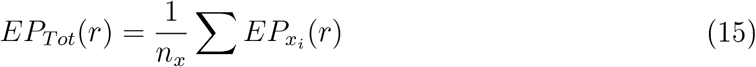

Where *EP*_*Tot*_(*r*) represents the overall behavior of the electric potential in the filament model as a function of distance away from the filament surface, *n*_*x*_ = 15 is the number of equally-spaced, perpendicular planes, and 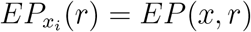 is the angular average electric potential obtained for each plane. Furthermore, the electrical double layer (EDL) thickness can be assessed from the computational results, thus allowing the calculation of the total current via a double integral method based on the setup employed for the average calculations described earlier. For instance, the total ionic current *I* in the EDL at the location *x* = *X*_0_ can be calculated using the double integral of the current density as a function of distance and angle

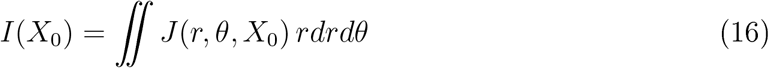

To implement this formulation, the values of current density obtained as a function of distance and angle from the plane located at *x* = *X*_0_, were exported as a table and rearranged using Matlab to calculate the double integral using the Trapezoidal method. Once the total current for all planes is obtained, the resistance was calculated using Ohm’s law as *R* = *V*_*sys*_*/I*_*T*_ where *V*_*sys*_ is the potential differential applied on the system, and *I*_*T*_ is the average total current obtained from the fifteen evaluation regions.

## Notes

### Competing Interest Statement

The authors have declared no competing interest.

